# Characterizing microplastic ingestion, transformation, and excretion in insects using fluorescent plastics

**DOI:** 10.1101/2022.11.11.516210

**Authors:** Marshall W. Ritchie, Alexandra Cheslock, Madelaine P.T. Bourdages, Bonnie M. Hamilton, Jennifer F. Provencher, Jane E. Allison, Heath A. MacMillan

## Abstract

Plastic pollution is a growing threat to our natural environment. Plastic waste/pollution results from high emissions of both macro (> 5 mm) and microplastics (MPs; < 5 mm) as well as environmental fractioning of macroplastics into microplastics. Microplastics have been shown to have a range of negative impacts on biota. Harmonized methods to accurately measure and count MPs from animal samples are limited, but what methods exist are not ideal for a controlled laboratory environment where plastic ingestion, transformation, and elimination can be quantified and related to molecular, physiological, and organismal traits. Here we propose a complete method for isolating and characterizing fluorescent MPs by combining several previously reported approaches into one comprehensive workflow. We combine tissue dissection, organic material digestion, sample filtering, and automated imaging techniques to show how fluorescently-labelled MPs provided to animals (e.g. in their diet) in a laboratory setting can be isolated, identified, and quantified. As a proof of concept, we fed crickets (*Gryllodes sigillatus*) a diet of 2.5% (w/w) fluorescently-labelled plastics and isolated and characterized plastic particles within the gut and frass.

## Introduction

Plastics are common polymers with many practical uses in our daily lives, such as clothing, electronic parts, and insulation, due to their durability, low density, and strength (Awasthi et al., 2017; Li et al., 2016). Plastics began to be mass produced in the 1950s and have since become a common form of mismanaged waste (Andrady and Neal, 2009). Plastic waste comes from many sources, such as post-consumer products, industrial uses, and improper recycling systems and practices (Barnes et al., 2009; Li et al., 2016; Wong et al., 2015). All these sources of waste ultimately enter natural environments, and approximately seven decades of plastic accumulation (approximately 4900 megatons; Geyer et al., 2017) in our environments are now having unintended consequences (Borrelle et al., 2020; Jambeck et al., 2015). Complete plastic degradation can take centuries, and the time required for breakdown depends on the plastic’s physical and chemical properties and the environment (Barnes et al., 2009). Fuller and Gautam (2016) observed that plastic contamination within the soil from industrialized areas was as high as 0.3-6.7% (w/w) plastic. Regardless of the timeline, large plastics entering the environment will ultimately break down into microplastics (MPs; plastics <5 mm in size; (Barnes et al., 2009; de Souza Machado et al., 2018). Microplastics have been shown to have a wide range of adverse effects in biota (Bucci et al., 2020). While there has been an increase in studies of microplastics, harmonization among commonly used or accepted methods is lacking (van Mourik et al., 2021).

Identifying MPs in abiotic and biotic samples requires controlled extraction and isolation steps and specialized equipment. Chemical digestions are often used (e.g., potassium hydroxide (KOH) to remove organic material as a curial first step for the identification process (Lusher et al., 2017). On the one hand, Fourier Transformed Infrared (FTIR), Raman spectroscopy (Raman), and Pyrolysis–gas chromatography-mass spectrometry (Py-GC/MS) have all become common approaches for identification (Käppler et al., 2016; Primpke et al., 2020; Shruti et al., 2022; Xu et al., 2019). These tools are now considered a requirement to identify MPs collected from the field, and knowledge of the types of plastic present in natural environments is essential to understanding their effects. On the other hand, laboratory-grade MPs (pre-identified by the manufacturer) used in experiments do not require this identification step, as the specific plastics are known and selected by the experimenter. Importantly, regardless of where the samples were obtained, it is critical MPs are properly filtered and isolated from the original source media.

Following sample digestion, sample filtering and imaging have been successfully combined with classification methods to understand MP characteristics (such as polymer type, MP fragment or fibre, and damage). These consist of effectively filtering MPs from environmental samples and applying deep learning programs to count and analyze the plastics (Lorenzo-Navarro et al., 2020; Lorenzo-Navarro et al., 2021). These methods work well when other materials in the sample (other than the plastics) can be removed. More freely accessible programs (such as ImageJ) have also been used to count and analyze MPs particles (Chen et al., 2021; Mukhanov et al., 2019; Park et al., 2022; Prata et al., 2019). Filtering and imaging approaches allow for more precise measurements of plastic uptake, elimination, and toxicological effects and the ability to characterize the effects of organisms on the plastics themselves (such as processing, degrading, and retaining).

Fluorescent dyes have also been used to examine MPs that have been isolated from environmental and organism samples (Chen et al., 2021; Maes et al., 2017; Prata et al., 2019). Using fluorescence allows for easy identification of plastics within samples, and the most common fluorescent dye used for this purpose is Nile red (Park et al., 2022; Shruti et al., 2022). When combined with sample filtration techniques, fluorescent dyes can allow for clear visualization and characterization of small MP particles within the low micron range (Maes et al., 2017; Park et al., 2022; Shruti et al., 2022) that may otherwise be missed. Commercially available MPs with a fluorescent label bound to them during manufacturing have also been used to track MPs in the digestive tract. Which then can be used analyze MP effects on organisms under laboratory conditions in terrestrial and aquatic species (Al-Jaibachi et al., 2018; Dawson et al., 2018; Fudlosid et al., 2022; Hasegawa and Nakaoka, 2021; Mazurais et al., 2015; Watts et al., 2014). Despite the various methods available, simple and reproducible methods have yet to coalesce for tracking, quantifying, and characterizing plastic particles within animal samples (Lusher et al., 2017).

Here, we outline a complete method to extract, filter, and capture images of MPs from animal samples. By combining existing knowledge of tissue dissection and digestion techniques, plastic filtering, and the quantification of fluorescent MPs within images, we show how to measure MP abundance and size in samples obtained from an animal model. This method is best used within a laboratory setting where levels of MP contamination and exposure to MPs are predetermined. Our approach avoids expensive software and equipment that can be challenging for many researchers to acquire and substantially reduces the time required to count MPs in samples manually. We suggest that this method quantifies MPs ingested and transformed by model and non-model animal species. By enhancing our ability to test the effects of MPs on animals and how animals may transform ingested plastics of different polymer types, sizes, and forms. As a proof-of-concept, we used this method to quantify the MP content of the gastrointestinal tract dissected from crickets fed MPs according to a standardized feeding protocol and track the excretion of MPs in cricket frass throughout their development.

## Methods

### Cricket rearing

All crickets (*Gryllodes sigillatus*) were reared from eggs obtained from Entomo Farms (Norwood, ON, Canada; an industrial partner raising crickets for human consumption). Freshly laid eggs were placed in an incubator at 32°C and held at approximately 40% relative humidity (RH) with a 14:10 Light: Dark (L:D) light cycle. Once the eggs hatched, cricket nymphs were randomly assigned to bins (32 × 28 × 19 cm). Juvenile crickets were given control food (Entomo farms cricket food mix containing a mixture of corn, soybean, herring, and hog meal) with unlimited access to water and shelter within these bins. Crickets were allowed to mature for one week to ensure any unexplained early-life mortality occurred before the crickets were assigned to treatment groups. Immediately after one week, the crickets were assigned to new bins (32 × 28 × 19 cm) and exposed to either the control diet or the same base diet containing 2.5% w/w of low-density Fluorescent Blue Polyethylene Microspheres ~100 μm (Cospheric, Fluorescent Blue Polyethylene Microspheres 1.13 g/cc 90-106 μm). At one week, forty juvenile crickets of unknown sex were placed in bins and held in their bins until reaching the age of 7-8 weeks. The bins contained an egg carton for shelter and an unlimited amount of food and water according to their feed diets, which were replenished each week when cleaned. All frass (insect digestive excrement) was collected during the cleaning, then frozen at −20°C for later analysis.

A stepwise protocol of the dissection, filtration, and image analysis steps that follow is included in the supplementary material.

### Gut dissection

Once crickets had finished their allotted exposure time and reached an age of seven weeks, they were dissected under the view of a dissection microscope (Zeiss, Stemi 508). All animals were weighed and anesthetized briefly with CO_2_ (15 s), placed in a petri dish filled with distilled water, and then vigorously dragged through the water to remove any MPs stuck to the cuticle. A visual inspection was done to look for any remaining plastics on the external surface of the animal using the dissection microscope. If clear, the washed crickets were then transferred to a second petri dish containing cricket saline (NaCl, CaCl_2_ (dihydrate), NaHCO_3_, Glucose, KCl, (recipe as in Whitaker et al., 2014). The crickets were carefully dissected from the dorsal side in saline under a dissecting microscope, focusing on keeping the entire digestive tract undamaged. The Malpighian tubules and tracheal system the were removed first. The esophagus of the cricket was then pinched and cut with forceps and pulled towards the cricket’s abdomen while gently teasing the digestive tract out of the body. The second pair of forceps were then used to pinch and cut the hindgut at the posterior end of the rectum before the entire gut was gently lifted out of the body. The cricket gut, once removed, was placed on a dry petri dish, where it was segmented into the foregut, midgut, and hindgut (anatomy and sections as in Figure 1). The segmenting was done dry (on a dish without saline) to ensure all MPs within the gut were left inside each region and were not washed out by saline. Early trials suggested that the proventriculus was resistant to KOH digestion when it remained intact, so this organ was torn open with forceps before sampling. The gut regions were blotted gently against a piece of paper towel to remove any extra saline stuck in the gut region. Each region was weighed and frozen at −20 °C for future use.

**Figure 1:**
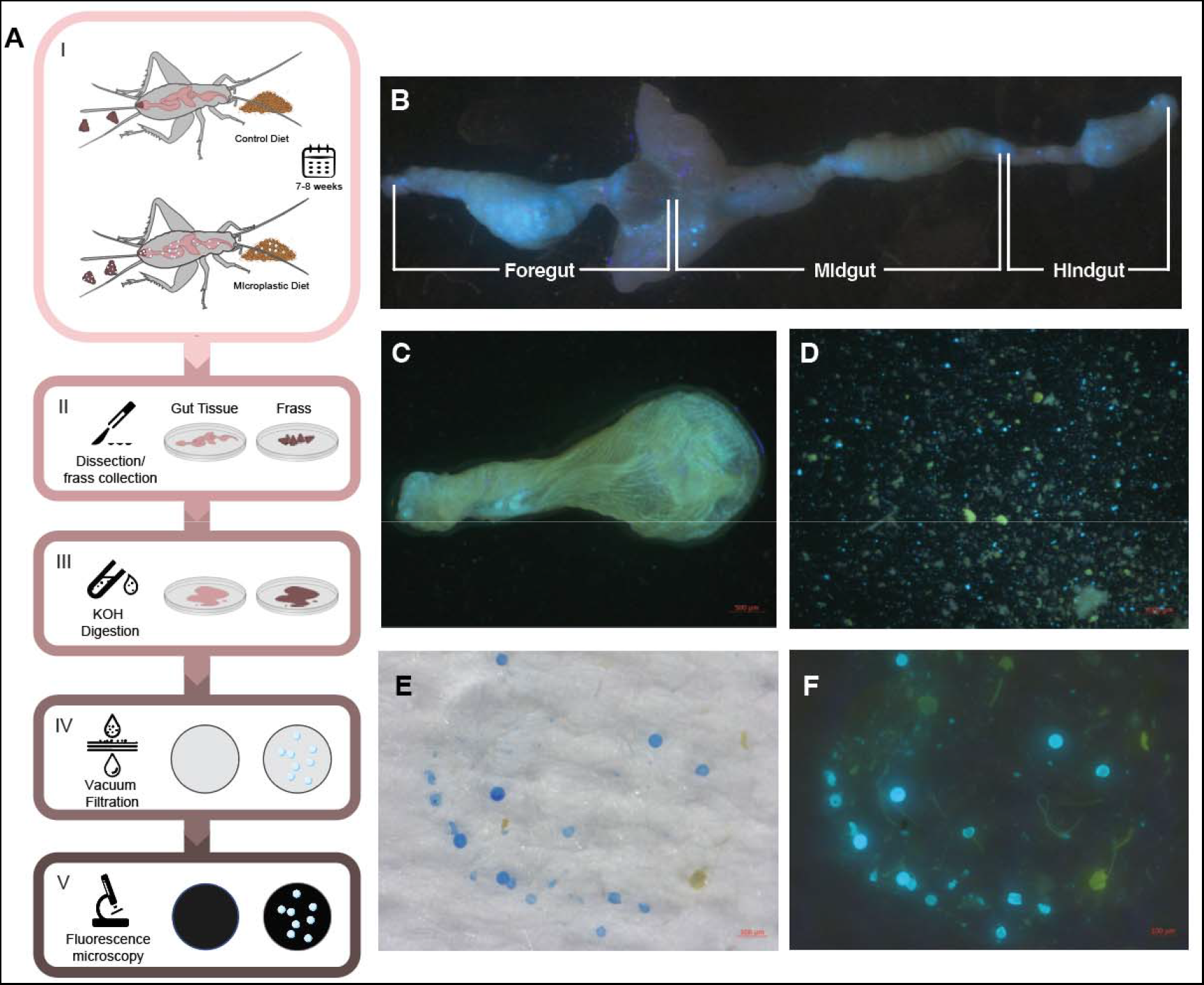
Graphical illustration of the workflow for isolating plastics from crickets (A). An intact digestive tract of an adult house cricket, *Gryllodes sigillatus* containing fluorescent plastic (B). Optionally, gut tissues can be cleared of their plastic contents. Using this approach the crop was cleared of its contents (C), and the contents from the crop were imaged without needing tissue digestion with KOH (D). Images of isolated plastics on the filters without fluorescence excitation (E) and the same filter with plastics fluorescing (F).

### Flushing of gut contents (optional)

In a subset of samples, we tested an alternative flushing approach to isolating plastics from the gut, which may be useful in some experiments (e.g. when isolating gut tissues from plastics for downstream biochemistry or molecular biology). Once the desired gut region was dissected from the specimen (as described above), the tissue was transferred to a clean petri dish with saline (enough to submerge the sample and allow an excess of flushing through the tissue). Before flushing, the other end of the gut region was cut with micro scissors to ensure the exit of the gut contents. To separate tissue from the contents, a 26-gauge needle with a blunted tip (cut) was used to avoid unnecessarily piercing tissue during flushing. Forceps were used to hold the tissue while the syringe (1ml) and blunted tip (pre-loaded with saline) were inserted into the lumen of the gut. Saline was then released from the syringe to create pressure and flush the gut contents from the lumen until no observable plastic (using a dissecting microscope) could be seen. Gut contents may be saved if desired by collecting and storing the saline used during the flushing process. Note that extra care is required for thin tissues (such as the midgut).

### Microplastic isolation and imaging

Intact gut regions that had not been flushed of contents from each group were removed from the freezer and thawed at room temperature. The gut regions were then digested in 10% KOH with different volumes used for the different gut regions based on earlier trials (200 μL for the foregut, 250 μL for the midgut, and 150 μL for the hindgut). Samples were held at 60°C for two days to digest, as described by Lusher et al. (2017). After two days, the samples were briefly vortexed before a subsample of 5 μL of the liquid containing the digested gut region was placed on a 1 μm pore-size glass fibre filter (Sigma, APFB04700) on top of a vacuum pump assembly (Sigma, Z290408-1EA). The subsamples were filtered using a hand pump (Fisher, S12932) attached to the vacuum pump assembly. After filtering, the filter paper and the MPs trapped on it were moved to a petri dish and photographed under a dissecting microscope (Zeiss, Stemi 508 with an axiocam 105 colour). The dissecting microscope was fitted with a 450 nm long-pass emission filter and 400-415 nm excitation light source (Nightsea LLC, Lexington, United States, Stereo Microscope Fluorescence Adapter).

### Frass digestion and imaging

The frass frozen from group reared crickets assigned to feed on the 2.5% w/w plastic diet was removed from the freezer and allowed to thaw completely. Once thawed, 15 pseudo-randomly selected individual frass pellets were isolated using forceps and individually placed in 0.6 mL microcentrifuge tubes. These frass samples were then digested with 15 μL of 10% KOH and left for two days, sealed at 60°C to ensure complete digestion of the frass. Finally, the frass samples were filtered and photographed using the same procedure as the gut regions. This was repeated for the six frass collection periods during cricket development.

### Image analysis

Images of fluorescent plastic particles were analyzed using ImageJ (FIJI version, 2.3.051; java, 1.8.0_172[64-bit]). To eliminate counting bias and reduce analysis time, we opted to use ImageJ’s predefined threshold ranges to count and measure the area of every plastic particle from the samples. Particles were removed from the dataset if they had areas larger than 20,000 μm^2^, as this was just over double the area of the largest beads identified in beads from the manufacturer bottle. Using the 20,000 μm^2^ cut-offs, 8 out of 2,557 captured samples (~0.3%) were removed from the analysis. An informal analysis of these images confirmed that these images contained particles that were too close together and thus resulted in an implausible particle area. Of the filter images, 7 (~8%) could also not be processed as there was too much plastic to accurately determine the area of individual pieces or plastics that were too small and could not be seen using this camera. No fluorescence signal was detected in any control samples (from crickets, not fed plastics).

To get accurate and unbiased counts of the plastics within each photo and to determine their sizes, we adapted a protocol from (Labno 2014). Our final image analysis method was as follows: The image was opened in ImageJ, the scale bar was measured using the straight-line tool, and the image scale was updated using the set scale function in ImageJ. Using the RGB stack function, the image was split into stacks by colour. The red stack was deleted as it was noted to create noise that impaired particle counting and measuring in the later steps. The Z project function was then used to combine the remaining two stacks of blue and green using the Average Intensity setting. This created a new combined image that was then adjusted using an auto threshold with either the Intermodes or Renyientropy option depending on light saturation and the amount of plastic present within the image (Cincotti et al., 2020). For images that contained many plastics in proximity, the Intermodes threshold was used; when images contained fewer particles or mainly small but well-separated particles, Renyientropy was used. The image was then converted to mask (binary) to create a white background with the fluorescent plastics appearing black. The watershed function was then run to reduce counting errors caused by plastic particles directly next to each other in the image. After these steps, the analyze particle function was used. The infinity (or size exclusion) was set to 5 units within the analyze particle function (meaning particles of a size less than 5 pixels were ignored). This step was completed to ensure any artificial particles created from the auto thresholding and watershed functions were removed from the final dataset but has the limitation of ignoring real particles at the limits of camera resolution. The circularity was set to 0-1.00 to capture any particles bigger than 5 pixels, regardless of shape. This function outputted the number of particles and the area in μm^2^ for each particle identified in the image.

### Comparison of automated counting vs manual counting

To determine the strengths and weaknesses of the above automated approach, we also validated this method against counts produced by impartial experimenters. Using the sample() function in R, 15 images were pseudo-randomly selected for blind manual counting from three independent researchers. The researchers were asked to count every visible blue fluorescent particle observed within each image without manipulating the image. These manual counts were then averaged and compared to the counts generated by the automated ImageJ process to validate the approach. The infinity was set to 5 to 1 (5 being what the frass measurements were set to) as a comparison to the researcher’s manual counts.

### Data analysis

All statistical analyses and plotting were completed in R version 4.1.2 (Foundation, 2022). The generalized linear model function (glm()), followed by a type III ANOVA (using the car package), was used to test for an effect of age (week of frass collection) on both the total area and the number of particles found within the frass pellets.

## Results and Discussion

The method outlined in this paper is an easy and reproducible workflow to measure surface area and count fluorescent MPs. Our approach combines and builds upon existing but fragmented techniques that have been previously described within the MP research community (Erni-Cassola et al., 2017; Labno, 2014; Thiele et al., 2019). The workflow described here combines these methods into a new straightforward, and effective workflow (Figure 1A). The method works by combining administering plastics to an organism, dissecting key regions within the organism, digesting the organic tissue to leave only plastics, filtering the plastics using widely available equipment and imaging, and automating characterization of those plastics with widely accessible software.

To demonstrate the utility of our approach, we used an insect model and tracked ingestion, passage through, and excretion from the digestive system. This approach is generally inexpensive and highly reproducible, and we believe it can be of great use to researchers interested in the effects of the animal digestive system on ingested plastics (transformation), tracking dosage of ingested plastics, and characterizing the effects of a given dose of plastic on animal physiology and fitness.

Fudlosid et al. (2022) found that crickets would ingest the same MP beads as those used here and noted that plastic was excreted, but the amount of plastic present in the gut and frass was not quantified. We first examined the segmenting and digestion of the foregut, midgut, and hindgut (Figure 1B) in the crickets to measure whether plastic was present and, if so, how much. We first isolated the plastic content of the gut regions from the gut tissues (e.g. crop; Figure 1C) by clearing them using a syringe and collecting the contents in a petri dish (Figure 1E). We found that clearing the gut contents allowed for good preservation of the tissue (e.g. for other downstream analyses) and allowed the contents to be almost entirely flushed out. However, to only characterize plastic content in a more high-throughput manner, we opted for a digestion approach.

Using the methods outlined above, we were able to segment the gut into its three major regions and effectively isolate plastic from each of the regions. This allowed us to capture clear images of the MP particles trapped within the 1 μm pore filters for quantification (Figure 2A). While we are confident in this approach, it is not without some limitations. Common errors in automated particle detection include missing smaller particles (false negatives), creating artificial particles through image processing (false positives), and artificially enhancing particle sizes (Bao et al., 2006; Erni-Cassola et al., 2017; Guan et al., 2018; Park et al., 2022). Each of these has consequences for hypothesis testing, such as under- or overestimating the number of particles in a sample and thereby under- or overestimating the total quantity of plastics and their size. These limitations can be seen when particles are overlapping and occasionally interpreted as a single particle (Blue circle; Figure 2B, C). We also found that particles from gut and frass samples were much smaller than originally anticipated, and the smallest particles could not be detected using the automatic thresholds because they occupied fewer pixels than our predetermined threshold (px = 5) (Red circle; Figure 2B, C). Capturing small MPs with automated thresholds has been similarly challenging for others (Mateos-Cárdenas et al., 2020), as particles under 20 μm were not detected (Erni-Cassola et al., 2017). In our case, we accounted for conjoined particles by first excluding impossibly large particles (possible because we controlled the size of plastics entering the animal), allowing us to count and measure most particles automatically (Figure 2D). The threshold setting for the ImageJ counts was subsequently lowered from 5 to 1 px to observe how threshold choice impacted counts. As the threshold was lowered, the number of detected particles increased, but the accuracy of correctly identified particles also changed.

**Figure 2:**
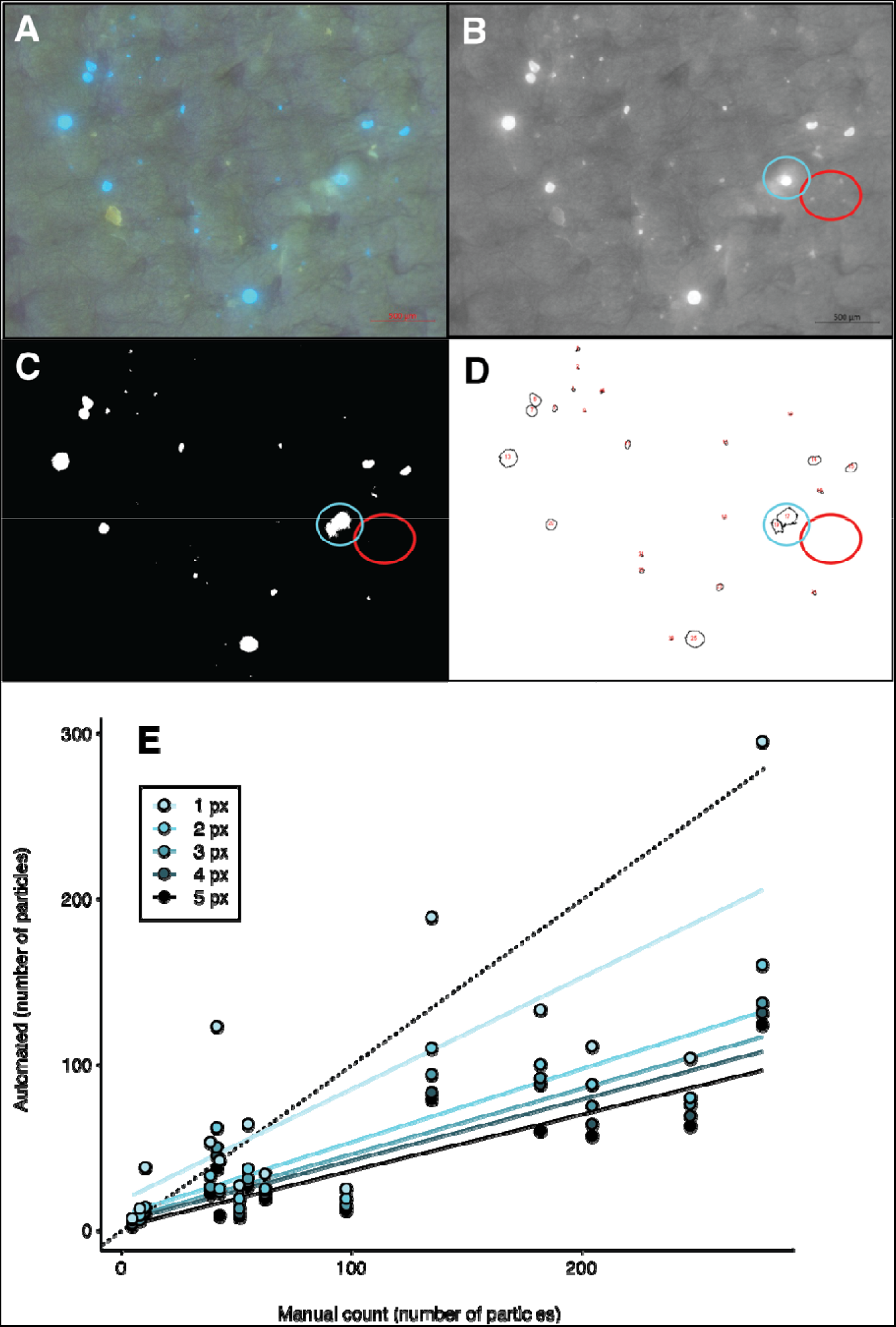
Example image of plastics from a cricket gut sample on a filter and image processing particle counting steps. Full-colour image (A) was converted to greyscale, and the red colour channel was removed (B). An auto threshold (using intermodes for this image) was used to convert the greyscale image to binary (C) and analyze the particle’s function, then highlight all particles analyzed (D). The blue circle highlights a particle whose area is exaggerated by ImageJ. The red circle highlights small visible particles that were not detected using this automated approach because they were smaller than our predefined threshold of 5 pixels. ImageJ thresholds (1-5 px) counts were plotted against average researcher counts using the same 15 images (E). The black dashed line shows a slope of 1.

To test the accuracy of the automated image analysis method, we had three researchers count particles in images with sample information redacted from the images. We found that the researchers identified more particles in the images visually than were detected using the automated approach, especially when images contain many plastic particles (Figure 2E). One likely reason for such a difference in manual and automated counts is that our predetermined ImageJ detection limit was removing small particles that were deemed real by the experimenters. To test this, we re-ran the same analysis with different threshold settings and found that lowering the setting did bring the slopes closer together, but the error within the detection increased. This was especially apparent at a threshold of 1 px, where any visible particle was counted. Acknowledging the limitations of the automated approach described above, we note that researchers generally did not arrive at the same counts as each other (Table S1). Thus, using automated counting software such as ImageJ to analyze particles removes human errors resulting from manual counting. We argue that the benefits of an automated approach outweigh these limitations as it will always produce the same counts and area from the same data when using the same settings (Park et al., 2022). Therefore, while this can introduce systematic errors that can and should be tested for and reported, the results remain more reproducible and objective than those from manual counts.

To demonstrate how our method can be applied to track plastic feeding and transformation, we fed crickets microplastics throughout their development and measured plastics in frass samples collected at one-week intervals. We expected that crickets would consume a greater quantity of plastics as crickets developed, and indeed there was a significant increase (*p*=0.003) in the total quantity of plastic (quantified as the total area of plastic within the images) as crickets aged and grew larger, which generally stayed consistent after the crickets completed their final moult (at approximately 5 weeks of age; Figure 3A). The same pattern (*p*= 0.004) was evident for the total number of plastic particles detected within the frass samples over developmental time (Figure 3B). These data suggest that age and body size may be important determinants of the quantity of plastic consumed, transformed, and excreted by insects and that *G. sigillatus* ingests the particles within their food indiscriminately during development.

**Figure 3:**
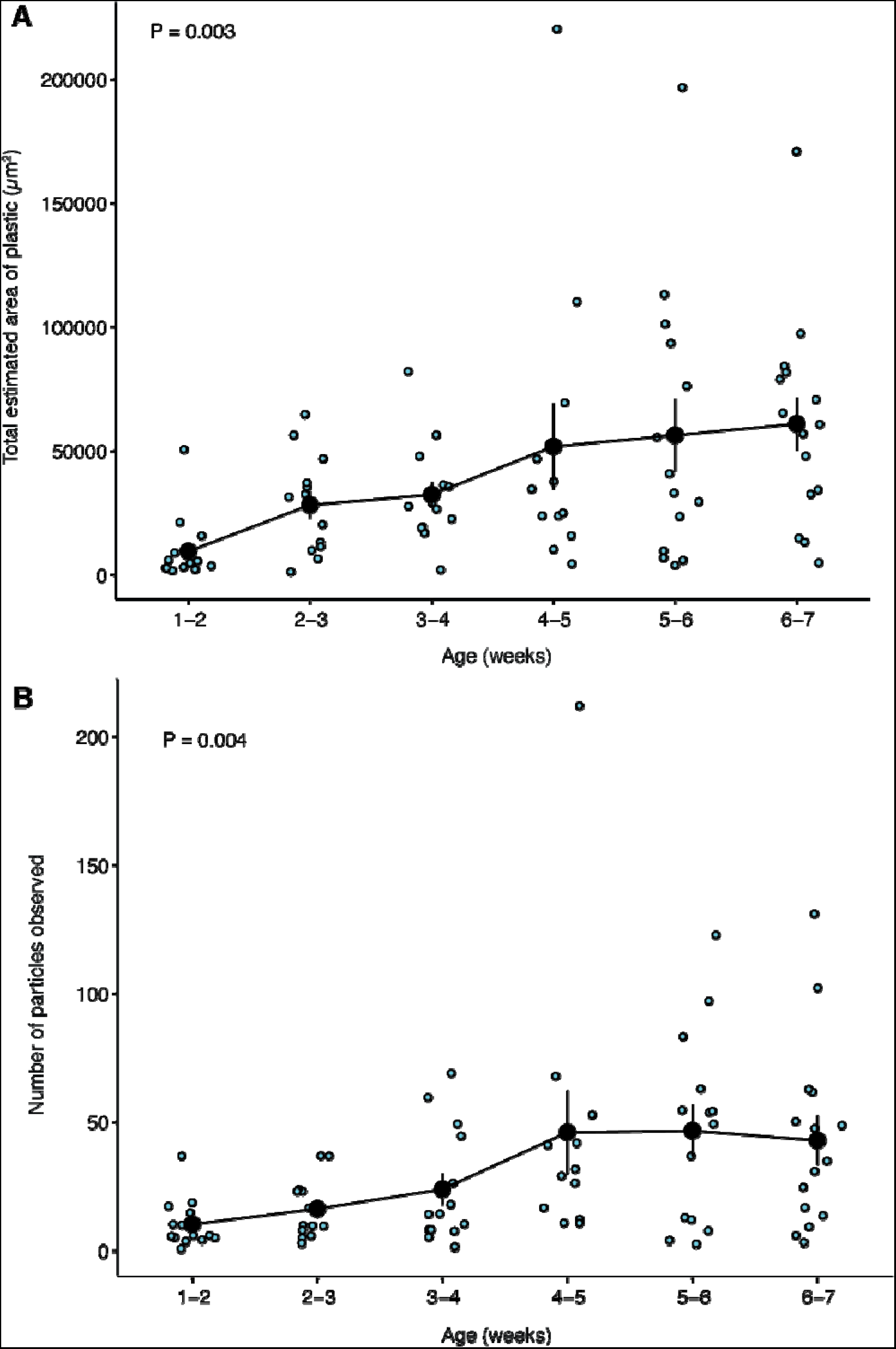
Fluorescent microplastics contained within the frass of *Gryllodes sigillatus* throughout ontogeny. (A) The total area of plastics within images of digested frass pallets collected throughout development. (B) The number of particles observed within each frass pellet during cricket development. Blue circles represent individual samples, and black circles represent the mean ± SE.

The methods outlined here for tissue dissection and digestion, as well as the filtering and isolating, and image analysis of microplastics, can be used and adapted as a standardized method to quantify MPs in a laboratory setting. Our approach uses widely available chemicals and equipment, and we argue that this method can be applied to a wide range of invertebrate and vertebrate species to produce accurate and reproducible counts of plastics in tissue and waste samples. Some of the limitations outlined here can be addressed with further refinement, and more powerful filtering and microscopy techniques capture would allow for measurement of any size of fluorescently labelled MPs and, perhaps, nanoplastics (NPs). We hope this method will be useful to those searching for a more standardized method to measure and characterize MPs as we seek to better understand how plastics are transferred and transformed, and what effects they can have on animal physiology and fitness.

## Acknowledgements

The authors would like to thank Mads Anderson for the support in discussions about the filtering process. The authors would also like to thank Matthew Muzzatti and Hannah Anderson for assisting with cricket care. We would also like to thank Entomo Farms for supplying cricket eggs during the project’s development.

## Conflict of Interest

The authors declare no conflicts of interest.

## Author Contributions

All authors conceived the study and designed the experiments. MR carried out the experiments. MR and HAM curated and analyzed the data and created visualizations. HAM and JFP provided resources and supervision. MR drafted the manuscript, and all authors edited the manuscript.

## Funding

This research was supported by funding from the Increasing Knowledge on Plastic Pollution Initiative from Environment and Climate Change Canada (project title: The fates and physiological consequences of plastics ingested by terrestrial arthropods) and a Natural Sciences and Engineering Research Council of Canada Discovery Grant (RGPIN-2018-05322) to HAM Equipment used in this study was acquired through support from the Canadian Foundation for Innovation and Ontario Research Fund (to HAM).

## Data availability

All data is provided as a supplementary file for review, and the same file will be included as supplementary material should the manuscript be accepted for publication.

## Notes

### Competing Interest Statement

The authors have declared no competing interest.

